# A novel approach for reliable differentiation of lymphatic endothelial progenitor cells in vitro

**DOI:** 10.1101/2025.09.23.677051

**Authors:** Inazio Arriola-Alvarez, Maria Rodriguez-Hidalgo, Hector Lafuente, Sonia Alonso-Martin, Ander Izeta

## Abstract

The lymphatic system plays critical roles in fluid homeostasis, immune regulation, and lipid transport, making its dysfunction a contributor to numerous pathological conditions. Lymphatic tissue engineering aims to develop therapeutic strategies for lymphatic regeneration and repair, depending on the availability of suitable cell sources for lymphatic endothelial cell (LEC) generation. Key cell sources explored for lymphatic tissue engineering include the dermis, bone marrow and stromal vascular fraction. To our knowledge, no existing differentiation protocols can accurately generate lymphatic endothelial progenitor cells (LEPCs). This study presents a novel *in vitro* differentiation protocol for generating LEPCs from stromal vascular fraction, dermis and bone marrow-derived mesenchymal stem cells (MSCs). The feasibility of using them as tissue sources was first evaluated leading to dermal cells exclusion due to low yield post-isolation, and highlighting the limited differentiation efficiency of bone marrow MSCs. However, the stromal vascular fraction emerged as the optimal source, exhibiting robust LEPCs differentiation and superior scalability. *In vitro* characterization confirmed that LEPCs maintain a lymphatic endothelial phenotype, exhibiting high migratory ability, robust angiogenic structure formation, and consistent expression of key lymphatic markers in both 2D and 3D cultures, as validated by RNA-Seq analysis. The protocol provides a consistent platform for studying lymphatic endothelial biology and potential applications in regenerative medicine and therapeutic angiogenesis.

## Introduction

The lymphatic system plays a critical role in maintaining tissue fluid homeostasis, immune response regulation, and lipid metabolism.^1^ Dysfunction in this system is implicated in a range of pathological conditions, including lymphedema, cancer metastasis, obesity, and neurodegenerative disorders.^2,3^ LECs form the structural foundation of the lymphatic system, constructing the vessel walls throughout the entire network. The system’s strongly hierarchical structure originates during embryogenesis, when a small subset of endothelial cells (ECs) in the cardinal vein differentiate into lymphatic lineage and begin expressing crucial markers like PDPN, LYVE1, and VEGFR3 around embryonic day 9.5 (E9.5).^4^ These cells, primarily guided by VEGF-C signaling via the VEGF-C/VEGFR3 pathway, develop into lymph sacs, which—along with non – venous origin LECs—form the mature lymphatic system.^5^ At this stage, additional factors such as IGF2, angiopoietins, HGF and FGF2 are also required for proper lymphatic development and maintenance.^6–8^ PDPN, a transmembrane glycoprotein, is critical for lymphatic vessel formation and function, playing roles in cell migration and maintaining lymphatic endothelial integrity.^9,10^ LYVE1 is involved in hyaluronan transport and serves as a key marker for LECs, indicating proper differentiation and functional potential of the cells.^11^ VEGFR3 is a primary receptor mediating VEGF-C signalling as described above, essential for lymphangiogenesis and the proliferation, migration, and survival of LECs.^12,13^

Recently, the limited but growing evidence supporting the importance of the lymphatic system in maintaining the tissue homeostasis and its implication on the diversity of diseases has gathered some interest, resulting in the development of few therapeutic approaches to stimulate lymphangiogenesis, i.e., the formation of new lymphatic vessels from preexisting ones.^14–16^

One of the strategies followed for this task is cell therapy. In this context, stem cells have gained significant interest for their potential in treating various diseases. These cells differ in potency, or their ability to differentiate into multiple cell types, with totipotent cells found only in embryos and adult stem cells, with lower potency –tissue specific–, residing in tissue niches to aid in repair and regeneration.^17^ Research has focused on harnessing adult stem cells for regenerative medicine, particularly MSCs. These are distinguished by their ability to form colonies, express specific surface markers, and maintain self-renewal while retaining differentiation potential, making them key to tissue engineering and therapeutic applications.^18–20^ Among these, bone marrow-derived (BM-MSCs), dermis-derived (D-MSCs), adipose tissue-derived (AD-MSCs) and umbilical cord-derived MSCs (UC-MSCs) have shown potential for lymphatic tissue engineering, each with distinct advantages.^21^ BM-MSCs, traditionally the most studied MSC type, have been explored for their ability to support lymphangiogenesis through paracrine signaling and direct differentiation.^22–25^ However, their invasive collection process and lower yield limit their practicality for large-scale applications.^26^ D-MSCs are a very invasive option but exhibit promising lymphangiogenic properties.^27,28^ Their role in supporting lymphatic endothelial cell differentiation has been noted, particularly through interactions with extracellular matrix components that resemble the native lymphatic niche.^29^ AD-MSCs offer a highly accessible and abundant cell source, with minimal donor morbidity.^30–32^ They have gained increasing attention for their potential in lymphatic tissue engineering due to their robust proliferation capacity, differentiation potential, and ability to secrete pro-lymphangiogenic factors, such as VEGF-C.^12,33,34^ UC-MSCs provide benefits, including easier and less invasive collection, enhanced proliferation capacity, and greater potency.^35,36^ Cord-blood endothelial cell colony-forming cells (ECFCs), have been recognized for their role in promoting lymphangiogenesis through the contribution of the lymphatic ECFC subpopulation they contain.^37,38^ Despite these promising attributes, tissue engineering has not put hardly any interest in their use for lymphatic tissue engineering.^21^ Overall, lymphatic tissue engineering lacks of adequate approaches to reliably differentiate MSCs into functional LECs *in vitro*.

Therefore, this study presents a novel differentiation protocol for deriving LEPCs from MSCs, focusing on dermis, bone marrow (BM) and adipose tissue-stromal vascular fraction (SVF) as the main cell sources. By leveraging key growth factors and signaling pathways involved in lymphatic embryogenesis and postnatal lymphangiogenesis, we have established a robust and reproducible method for generating functional LEPCs. Furthermore, in this work, we have proven that SVF-derived LEPCs display the best differentiation potential, and coupled with their ease of access and proliferation capacity, pose as a great alternative for LEPC differentiation in the clinical settings. This work not only provides a valuable tool for advancing lymphatic disease modeling and therapeutic applications but also highlights the broader potential of MSCs in lymphatic tissue engineering.

## Materials and methods

### Isolation of SVF, BM and dermis

SVF and BM were extracted from C57BL/6 mice (8 week old male and female). Briefly, adipose tissue was scraped off the inner part of the back skin. Then, it was minced into approximately 1 mm^2^ pieces, and incubated under 180 rpm agitation in a collagenase type IA solution (1.000 u/mL, C2674, Sigma) for approximately 25 minutes. Later, the digested adipose tissue solution was inactivated and centrifuged at 1.500 rpm for 5 minutes to separate into layers. The top two layers were removed and the bottom layer (containing the SVF’s cells) was washed twice with 2% fetal bovine serum (FBS, 10270-106, Gibco) in D-PBS (14190-094, Gibco), filtered at 100 µm and plated for culture. Besides, the femur, the tibia and the humerus were extracted from the mice, and the epiphysis of the bones were cut leaving an open wound. Afterwards, the bones were placed in a 0.5 mL tube with a small hole in the bottom. Then, these two were placed inside a 1.5 mL tube and centrifuged at 10.000 rpm for 10 seconds. The BM was flushed out into the 1.5 mL whilst the empty bones were retained in the smaller tubes. The BM pellet was washed twice again with 2% FBS and plated for culture. Skin specimens clean of blood vessels and adipose tissue were minced into 1-2 mm² fragments. Digestion was performed in 1 mg/ml collagenase XI at 37°C for ∼60 min until tissue fragments became pale and easily dispersible. Digestion was stopped with cold wash medium. After centrifugation (300g, 7 min, 4 °C), pellets were mechanically dissociated in wash medium via pipette grinding (2-3 min/repetition) until complete dissociation (milky suspension). Cells were filtered (40 μm), centrifuged, and resuspended in proliferation medium. Viability was assessed by trypan blue exclusion.

### Cell culture

Isolated SVF-derived MSCs were directly seeded onto the plastic plates with complete Mesencult medium (05513, StemCell) and cultured in a humidified incubator with a gas mixture of 5% CO_2_ at 37°C. Bone marrow MSCs were also cultured in complete Mesencult medium in a humidified incubator with a gas mixture of 1% O_2_, 5% CO_2_ at 37°C. MSCs were seeded at 10.000 cell/cm^2^ and cultured until passage 3. Once cells had reached 80% confluence, they were transferred to new dishes using 0.25% Trypsin-EDTA (25200-072, Gibco). Human dermal lymphatic endothelial cells (HDLEC, C-14021, PromoCell) were cultured according to the providers’ guidelines.

### Adipogenesis

MSCs isolated from SVF and BM were expanded to passage 2 using standard subculture techniques. At 90% confluence, cells were trypsinized and seeded at 10,000 cells/cm² in 12-well plates. Differentiation was induced using the MesenCult Adipogenic Differentiation Kit (05507, StemCell) under hypoxia (1% O₂, 5% CO₂, N₂-balanced) in a tri-gas incubator. The medium was refreshed every 3–4 days, with differentiation periods of 14 days (SVF-derived MSCs) or 7 days (BM-derived MSCs), as per the manufacturer’s guidelines. Post-differentiation, cells were fixed with Histofix (256462, Panreac AppliChem) for 10 min and stored in dPBS for subsequent staining.

### Osteogenesis

MSCs at passage 2 were seeded at 10,000 cells/cm² in 12-well plates and differentiated using the MesenCult Osteogenic Stimulatory Kit (05504, StemCell) upon reaching 90% confluence. Cells were cultured under hypoxia (1% O₂, 5% CO₂) with medium changes every 3-4 days for 21 days (SVF-derived MSCs) or 7 days (BM-derived MSCs), as per manufacturer’s guidelines. Post-differentiation, cells were fixed in Histofix for 10 min and stored in dPBS for subsequent staining.

### Histological analysis

To confirm adipogenesis, cells were stained with Oil Red O (O0625, Sigma). A stock solution (300 mg Oil Red O in 99 mL 99% isopropanol) was diluted 3:2 with deionized water to prepare the working solution, filtered (0.22 μm), and used within 2 h. Fixed cells were washed with dPBS, treated with 60% isopropanol (5 min) (59310, Sigma) then stained with Oil Red O (5 min). After rinsing, nuclei were counterstained with hematoxylin (1 min) and imaged using AxioObserver 7 microscope (Zeiss). To confirm osteogenesis, a 40 mM Alizarin Red S solution (A5533, Sigma) was prepared by dissolving 6.846 g in 400 mL deionized water, adjusting pH to 4.2 with NaOH (131687.1211, Panreac), and bringing to 500 mL final volume before 0.22 μm filtration. Fixed samples were washed with dPBS, air-dried, then stained with 1 mL/well Alizarin Red solution (15 min, RT). After gentle aspiration, samples were washed 3x with tap water (1 mL/well) while preserving mineralized deposits. Imaging was performed using an AxioObserver 7 microscope (Zeiss).

### Differentiation of isolated MSCs toward LEPCs

At passage 3, MSCs were transferred to new dishes using 0.25% Trypsin-EDTA, and seeded at 20.000 cells/cm^2^ with complete mouse endothelial cell medium (ECGM, M1168, Cell Biologics) supplemented with 50 ng/mL of VEGF-C (752-VC, RD Systems), CCBE1 (H00147372-P01, Abnova), angiopoietin-2 (7186-AN, Angiopoietin-2), IGF2 (792-MG, RD Systems), FGF2 (3339-FB, RD Systems) and angiopoietin-1 (9936-AN, RD Systems). The differentiation protocol was split into 3 stages: induction, differentiation and maturation. Induction was carried for 5 days and cells were cultured with ECGM supplemented with VEGF-C.

Differentiation followed the previous and lasted 7 days cultured in ECGM supplemented with VEGF-C, CCBE1 and angiopoietin-2. Maturation comprised an additional 7 days of culture with ECGM supplemented with VEGF-C, IGF2, FGF2 and angiopoietin-1. Cells were maintained under standard culture conditions (37 °C with 5% CO_2_).

### Immunofluorescence staining

Cultured LEPCs and MSCs were fixed with Hystofix for 15 minutes diluted 1:1 in D-PBS, blocked and permeabilized with 10% FBS and 0.3% Triton X-100 (T8787, Sigma) in D-PBS and immunostained. Briefly, cells were immunostained with: anti-mouse PDPN (127402, BioLegend), anti-mouse LYVE1 (103-PA50AG, Reliatech) and anti-mouse VEGFR3 (AF743, RD Systems) as the primary antibodies; and Alexa Fluor 647-conjugated anti-hamster (A21451, Invitrogen), Alexa Fluor 555-conjugated anti-rabbit (A31572, Molecular Probes) and Alexa Fluor 488-conjugated anti-goat (A11055, Molecular Probes) as secondary antibodies. Alexa Fluor 488-conjugated phalloidin (A12379, Invitrogen) was used to selectively stain f-actin. 4′, 6-Diamidino-2-Phenylindole (DAPI, D3571, Invitrogen) was used to stain nuclei (please refer to table S1 for antibody details). Immunofluorescence images were taken using Zeiss AxioObserver 7 and LSM900 confocal microscope (Zeiss).

### Flow cytometry analysis

The expression of PDPN, SCA-1, CD90.2, CD105, CD44, CD31 (PECAM1) and CD34 of the isolated and differentiated cells was detected using the APC-conjugated anti-mouse PDPN (127410, BioLegend), PE-Cy7-conjugated anti-mouse SCA-1 (558162, BioLegend), anti-mouse CD90.2 (553009, BD Biosciences), anti-mouse CD105 (14-1051-82, ThermoFisher), anti-mouse CD44 (553134, BD Biosciences), PE-conjugated anti-mouse CD31 (553373, BD Biosciences) and PE-conjugated anti-mouse CD34 (551387, BD Biosciences).

Immunoglobulin isotype controls were used to distinguish specific from non-specific bindings during flow cytometry (402012, Biolegend; 553930, BD Biosciences). Alexa Fluor 488-conjugated secondary antibody was used for the unconjugated CD90.2 and CD105 primary antibodies (A11006, Molecular Probes)(please refer to table S2 for antibody details). The expression levels were determined by the median fluorescence intensity (MFI), using FlowJo v10 software (BD Biosciences).

### RNA extraction and RT-qPCR

Total RNA was isolated using tri reagent solution (AM9738, Life Technologies). Reverse transcription was performed using the High-Capacity cDNA reverse transcription kit (4368814, AppliedBiosystems) according to manufacturer’s guidelines. Quantitative real-time PCR was performed in a CFX384 Thermal Cycler (Bio-Rad) using TaqMan Gene Expression Master Mix (4369016, ThermoFisher) according to the manufacturer’s guidelines and 10 ng of cDNA. *Gapdh* and *Tbp* were used as housekeeping genes (please refer to table S3 for probe details). Statistical analysis was performed based on the difference between the Ct value of the target gene and the Ct value of the housekeeping gene (2^-ΔCt^ values). Gene expression was represented using 2^-ΔΔCt^ values.

### Genotype stability over passages

To assess the stability of the gene expression over the passages, cells were plated at 10.000 cells/cm^2^ on 12-well plates and cultured at 37 °C until 80% confluence was reached. Then, cells were detached (using Trypsin-EDTA 0.25%), counted and a fraction of them were seeded (10.000 cells/cm^2^) while the rest were lysed for RNA extraction at each passage until passage 6.

### Scratch-wound assay

Cells (10.000 cells/cm^2^) were plated on 24-well tissue culture plates and allowed to reach confluence (3-4 days). Once confluent, the media was aspirated and the cell layers wounded using a 20 µL micropipette tip. Then, the plates were rinsed with D-PBS to remove detached cells; fed with cell medium, and monitored until the wound had fully closed. Photographs were taken every 24 hours and images were analyzed using the Wound Healing tool plugin for ImageJ.

### Tube formation assay *in vitro*

10 µL of Matrigel (3 mg/ml, 354234, Corning) were added to each well (µ-slide 15 well, Ibidi) and left to harden for approximately 30 minutes. Once the coating had cross-linked, 10.000 cells in 50 µL were seeded directly onto the Matrigel coated wells, incubated at 37 °C for 5 hours, stained for LIVE/DEAD and photographed using Zeiss AxioObserver 7 (Zeiss).

### Three-dimensional (3D) culture *in vitro*

1×10^5^ cells were embedded in 100µL of Matrigel (3 mg/ml, 354234, Corning), seeded in an 8-well culture slide (BD Falcon) and let to crosslink for approximately 30 minutes. Once the gel had cross-linked, culture media was added and the constructs were incubated at 37 °C for 10 days. After 10 days, bright field images were taken to assess the web formation (AxioObserver 7, Zeiss), cell viability by LIVE/DEAD staining (L3224, Invitrogen) and fixed for immunofluorescence staining using Hystofix as described before.

### RNA Sequencing

Total RNA was extracted using TRI reagent solution. RNA libraries were sequenced on an Illumina NextSeq2000, generating 20-30 million paired-end reads (100 bp, 50 bp x 2). RNA-Seq data processing and differential expression analysis were performed using the standardized nf-core/rnaseq (v3.15.0) and nf-core/differentialabundance (v1.5.0) pipelines to ensure reproducibility and best practices in computational workflows.^39^ Raw sequencing data were processed through nf-core/rnaseq. Low-quality bases filtering (Phred score < 15) using a sliding window approach and adapter trimming were performed using fastp v.0.23.4.^40^ Ribosomal RNA (rRNA) was removed using SortMeRNA v.4.3.6.^41^ Cleaned reads were aligned to the *Mus musculus* GRCm39 reference genome (ENSEMBL) using STAR aligner v.2.7.10a.^42^ Transcript abundance was quantified at the gene level with Salmon v.1.10.1.^43^ Count matrices generated by nf-core/rnaseq were analyzed for differential expression using nf-core/differentialabundance, which leverages DESeq2 (v1.44.0).^44^ The design matrix explicitly accounted for technical variability from sample replication batches while estimating absolute expression levels across conditions. *p*-values were adjusted with the Benjamini-Hochberg (BH) method to reduce the number of false positives. Differentially expressed genes were called with thresholds of |log2(fold change)| ≥ 1 and adjusted *p*-value (FDR) < 0.05.

## Results

### MSC isolation is consistently more efficient for SVF than BM or dermis

SVF and BM display better results when it comes to cell count after the extraction protocol (Sup. 1). Whole BM cell suspensions contained approx. 250×10^6^ cells (>90% viability), while SVF approx. 2×10^6^ cells (>90% viability), and the dermis around 5×10^5^ with really poor viability (∼50%). Regarding these results dermis was discarded and our focus relied into BM and SVF. When cultured in complete Mesencult Medium, SVF-derived MSCs showed higher proliferation rate in comparison to BM-derived MSCs, showing twice the accumulated cell proliferation per mouse for almost half the time of culture (Sup. 1B). Moreover, cell culture showed less morphological heterogeneity in SVF-derived MSCs than BM-derived MSCs (Sup. 1A.II and C.II). Finally, MSC lineage was confirmed for both tissue sources by (a) analysis of MSC surface markers through flow cytometry and (b) their differentiation potential for adipogenesis and osteogenesis (Sup. 1A.III-V and C.III-V). Both SVF- and BM-derived cells showed typical MSC marker expression patterns: SVF-derived MSCs expressed high levels of CD90.2 (90.48% ± 4.54), CD105 (51.13% ± 4.87), and SCA1 (92.48% ± 2.10), along with moderate CD44 expression (28.5% ± 25.17); BM-MSCs exhibited lower CD90.2 (9.06% ± 0.23) and CD44 (12.42% ± 11.57) but maintained strong CD105 (61.05% ± 0.78) and SCA1 (82.45% ± 1.77) expression. Importantly, both populations showed minimal endothelial marker presence (SVF: CD31 0.08% ± 0.02, CD34 0.09% ± 0.05; BM: CD31 0.68% ± 0.39, CD34 0.8% ± 0.3). Both cell populations successfully underwent adipogenic and osteogenic differentiation, as evidenced by distinct histological staining patterns. Adipogenic differentiation was confirmed through the accumulation of lipid droplets that stained positively with Oil Red O, while osteogenic differentiation was verified by Alizarin Red S staining of calcium phosphate deposits and the morphological presence of mineralized nodules.

### Flow cytometry analysis shows that SVF-derived MSCs outperform BM-derived MSCs in lymphangiogenic differentiation potential

Flow cytometry analysis of SVF- and BM-derived LEPCs showed a significant increase in the expression of PDPN for both cell types (fig. 2A-B); nonetheless, SVF-derived LEPCs showed a higher increase than the BM (97.4% ± 2.2 of PDPN-positive LEPCs for SVF vs. 76.2% ± 19.5 for BM) (fig. 2A.I and B.I). In addition, MFI analysis confirm a statistically significant change in SVF-derived LEPCs (5.5×10^5^ MFI for LEPC vs. 1×10^5^ MFI for MSC; *p*-value<0.01) (fig. 2A.II). MFI results for BM-derived LEPCs also showed a statistically significant increase in PDPN expression (2.5×10^4^ MFI for LEPC vs. 1×10^4^ MFI for MSC; *p*-value<0.05) (fig. 2B.II). Notably, SVF-derived LEPCs exhibited significantly greater upregulation of PDPN expression compared to their BM-derived counterparts, as demonstrated by both higher absolute expression levels and enhanced fluorescence intensity quantification. Given these pronounced differences, we prioritized comprehensive characterization of the SVF-derived LEPCs.

**Fig 1.**
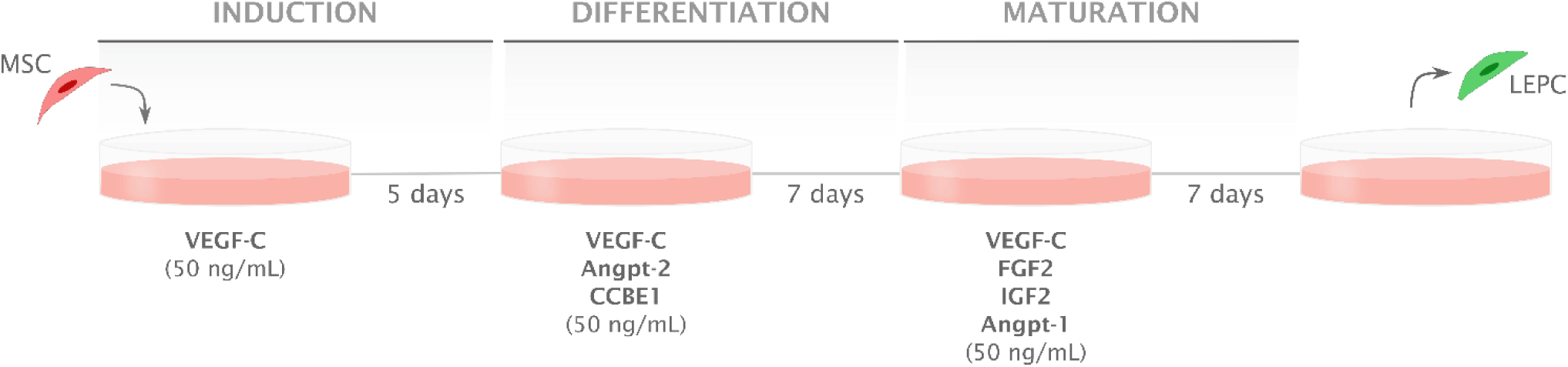
Differentiation protocol illustration. ***VEGF-C***: Vascular endothelial growth factor C. ***Angpt 2***: Angiopoietin 2. ***CCBE1***: Collagen and calcium binding EGF domains 1. ***FGF2***: Fibroblast growth factor 2. ***IGF2***: Insulin-like growth factor 2. ***Angpt 1***: Angiopoietin 1.

**Fig 2.**
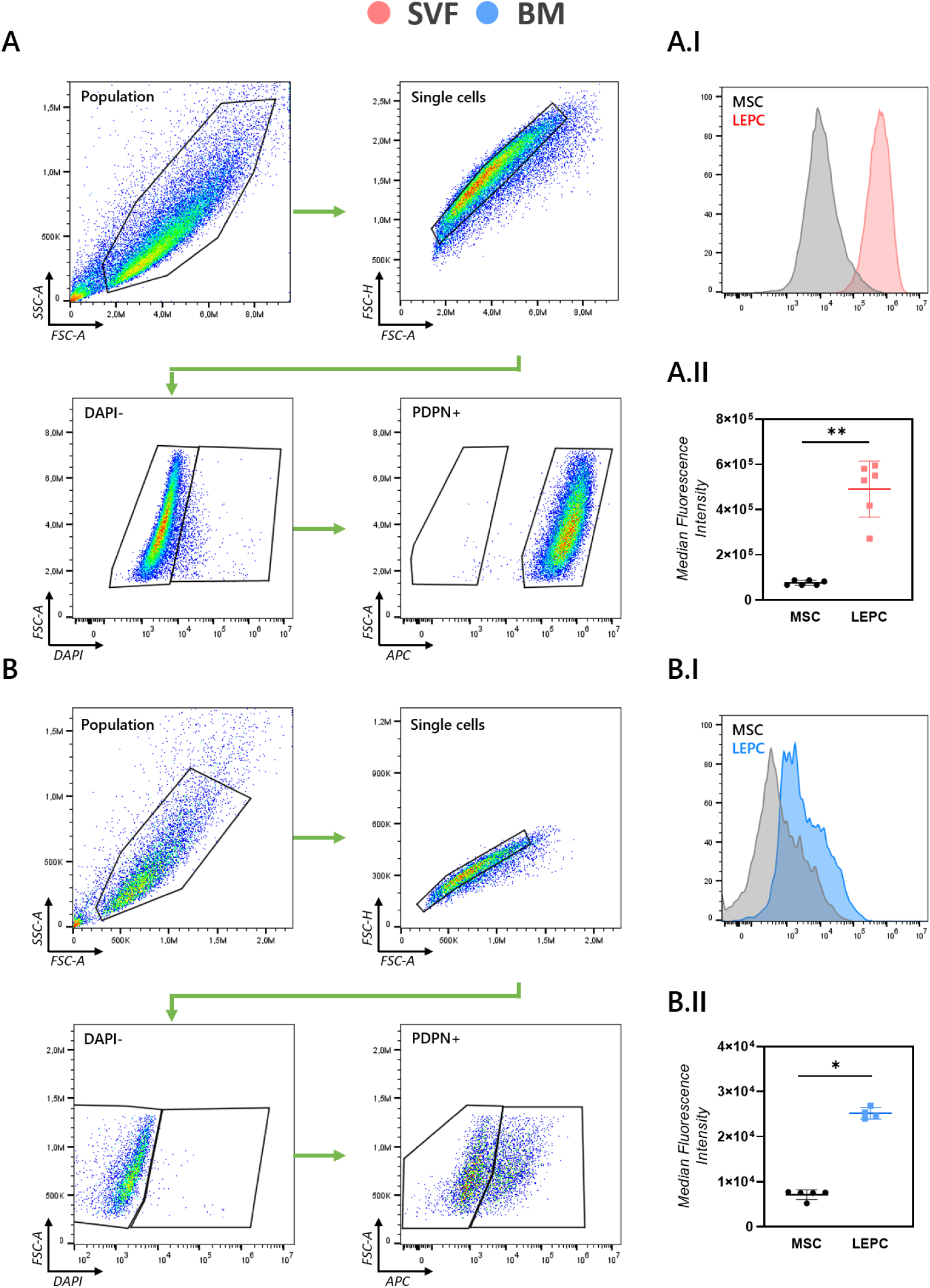
Stromal vascular fraction (SVF)-derived lymphatic endothelial progenitor cells (LEPCs) show better differentiation by flow cytometry analysis than bone marrow (BM)-derived LEPCs. (A) Flow cytometry workflow of the SVF-derived LEPCs from the first population until the PDPN-positive population. (A.I) PDPN expression histograms showing the expression level for SVF-LEPCs and -MSCs. (A.II) Median fluorescence intensity (MFI) of PDPN in SVF-LEPCs and –MSCs (mean ± SD). (B) Flow cytometry workflow of the BM-derived LEPCs from the first population until the PDPN-positive population (B.I) PDPN expression histograms showing the expression level for BM-LEPCs and -MSCs. (B.II) MFI of PDPN in BM-LEPCs and –MSCs (mean ± SD). Statistical significance: \**p*-value<0.5, \*\**p*-value<0.01. Mann-Whitney test. n=6 for SVF-LEPCs and n=5 for BM-LEPCs. Mean ± SD represented.

### LEPC lineage is confirmed by gene expression analysis and immunofluorescence staining of SVF-derived LEPC

Our novel differentiation protocol mimics the natural process of lymphatic embryogenesis and incorporates key molecular factors involved in adult lymphangiogenesis. By using VEGF-C, CCBE1, Ang1, Ang2, IGF2 and FGF2, the protocol demonstrated strong potential due to the complementary and synergistic roles these factors play in lymphatic vessel development and function^45^. RT-qPCR analysis confirmed the lymphatic commitment of SVF-derived LEPCs (Fig. 3A), with significant fold-changes observed in all critical lymphatic markers (*Pdpn, Lyve1, Vegfr3, Igf2, Vegf-c*), establishing their successful differentiation from MSC precursors (*p*-value<0.05). Immunofluorescence analysis of PDPN revealed a dramatic upregulation in SVF-derived LEPCs compared to their MSC progenitors, corroborating earlier findings (Fig. 3B). Quantitative assessment by ImageJ demonstrated a significant increase in MFI for SVF-derived LEPCs when normalized to nuclear counts within each sample (*p*-value<0.01).

**Fig 3.**
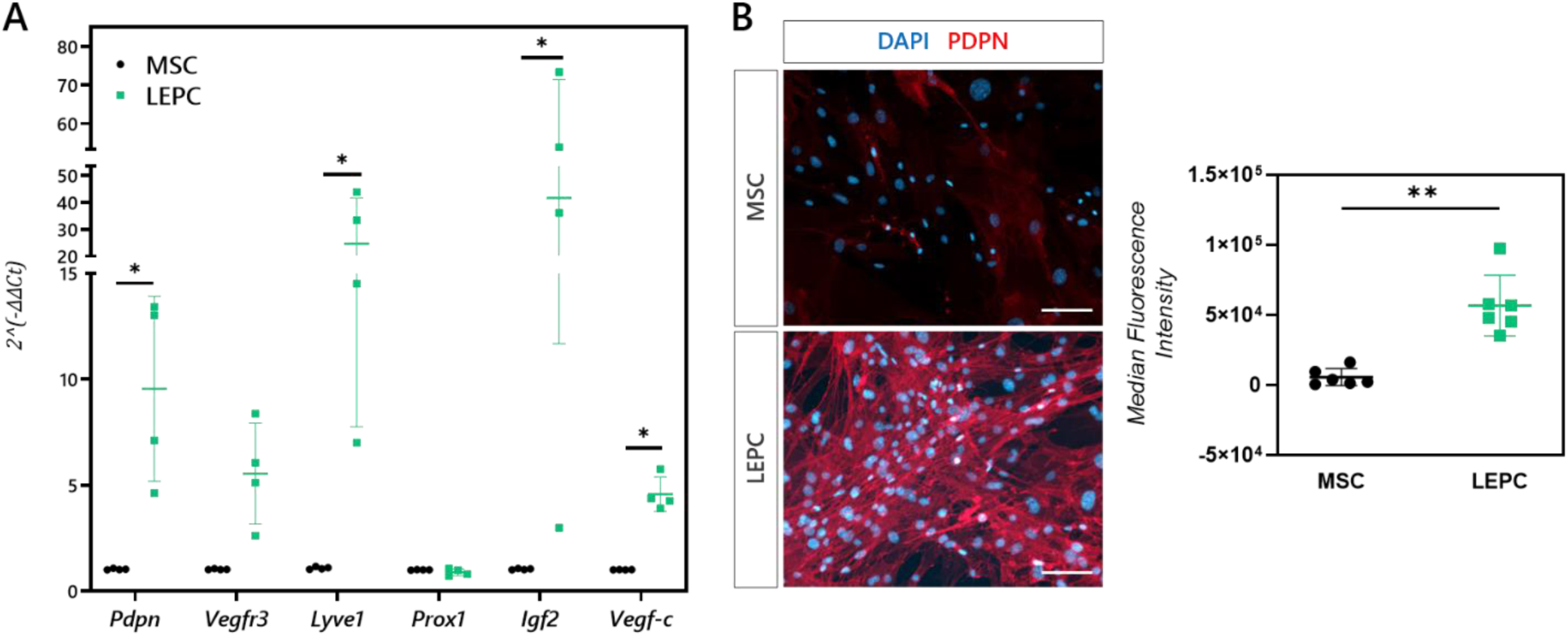
Characterization of lymphatic endothelial progenitor cells (LEPCs) shows correct differentiation by overexpression of lymphatic lineage specific markers. (A) RT-qPCR shows statistically significant overexpression of key lymphatic markers such as *Pdpn*, *Vegfr3*, *Lyve1*, *Vegf-c* and *Igf2*. n=4. MSC, mesenchymal stem cells. Mean ± SD. (B) Median fluorescence intensity analysis of immunofluorescence staining images shows statistically significant differential expression of PDPN between MSCs and LEPCs. Scale bar: 100 µm. n=6. Statistical significance: \**p*-value<0.05; \*\**p*-value<0.01. Mann-Whitney test.

### SVF-derived LEPCs show a reliable lymphatic phenotype *in vitro*

Our novel differentiation protocol was further characterized *in vitro* for genotype stability with 3 more passages after differentiation and analyzing the migratory phenotype via scratch-wound assay. The SVF-derived LEPCs showed little variation of gene expression of the key molecular markers of the lymphatic system for at least 1 passage after differentiation (P4) (fig. 4A). Moreover, SVF-derived LEPCs showed a very pro-migratory phenotype, closing the generated wound in less than 48 hours of culture (fig. 4B). Comparative analysis of cellular migration patterns revealed that our differentiated cells exhibited a migratory phenotype functionally equivalent to HDLECs. These control HDLECs served as our biological reference standard, with quantitative assessments demonstrating comparable migration rates between the two cell populations.

**Fig 4.**
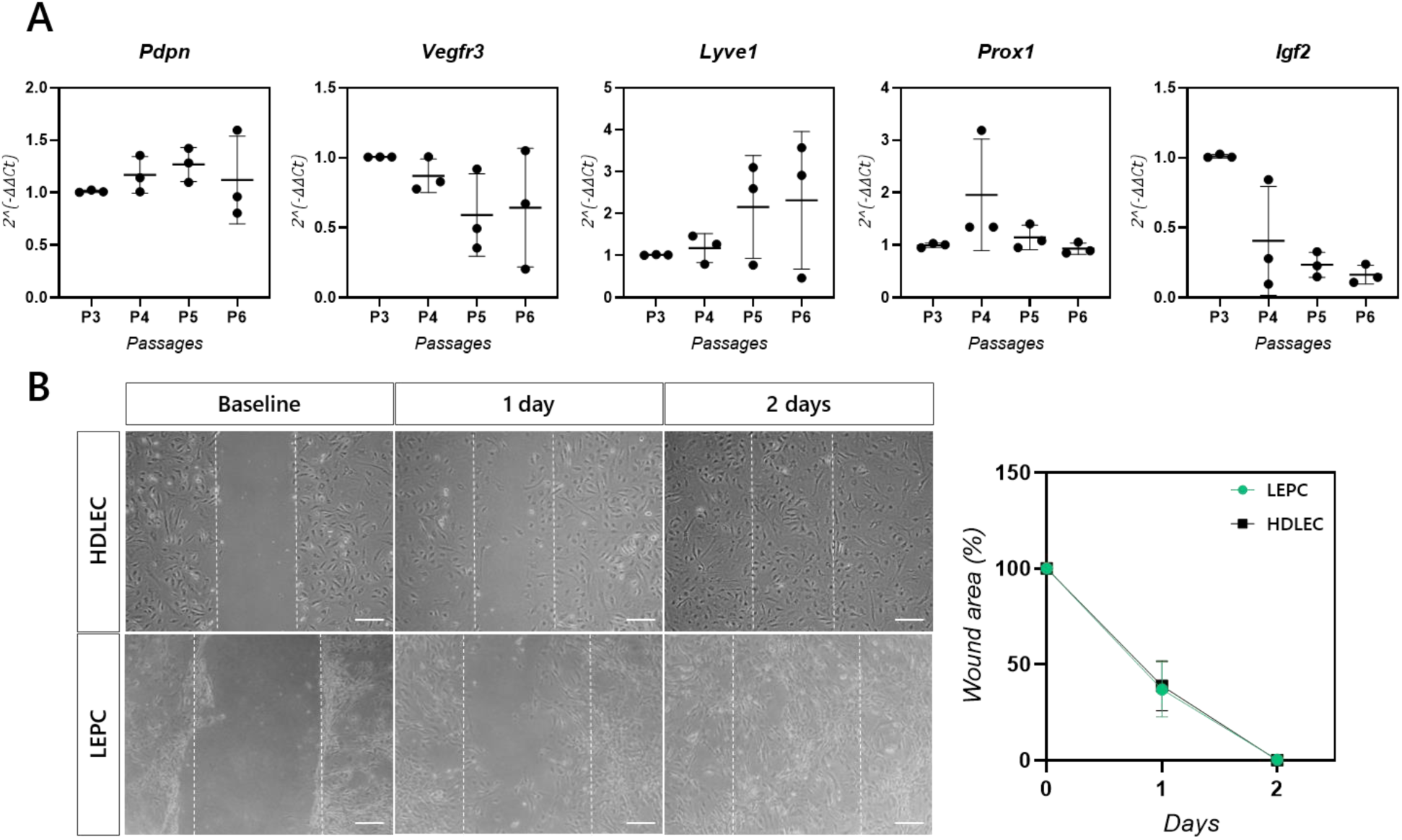
Stromal vascular fraction (SVF)-derived lymphatic endothelial progenitor cells (LEPCs) show a reliable lymphatic phenotype *in vitro.* (A) Genotype stability results of the key overexpressed genes over passages (P), assessed by RT-qPCR. n=3. (B) Scratch-wound assay of the SVF-derived LEPCs in comparison to human dermal lymphatic endothelial cells (HDLECs) along two days in culture. n=3. Scale bar: 200 µm. Mean ± SD. Mann-Whitney test.

### SVF-derived LEPCs outperform HDLECs in a biomimetic environment *in vitro*

In order to assess key functional characteristics of LEPCs, we assessed angiogenesis *in vitro* in 2D and 3D environments using Matrigel as the hydrogel matrix, and was compared with HDLECs as gold standards (control). We selected these cells as controls for several key reasons. First, they exhibit a well-defined lymphatic expression profile, with strong expression of established lymphatic markers.^46^ Second, they are readily accessible, being commercially available and widely referenced in existing studies (over 200 studies). Third, they are functionally validated, demonstrating characteristics consistent with LECs.^47^ Finally, we opted for human-derived cells over murine models to enhance clinical relevance, ensuring closer alignment with human physiology for translational applications.^48^ In a 2D manner, our SVF-derived LEPCs showed a similar capacity for capillary-like structure formation as the control, showing no statistically significant differences in the number of nodes, number of segments, number of loops and number of meshes (fig. 5A-B). Conversely, when embedded in a 3D Matrigel hydrogel, the SVF-derived LEPCs outperformed the control, showing significantly better tube-like formation (fig. 5C-E). Specifically, when cultured up to 10 days, while HDLECs showed a little tube-like formation or reorganization, remaining static, SVF-derived LEPCs showed a readily assembled web-like structure that matures into thick structures (fig. 5C-E). The average number of nodes, junctions, total number of meshes and master junctions for SVF-derived LEPCs greatly outperformed the dermal controls (fig. 5E). Finally, immunofluorescence staining of 3D cultured SVF-derived LEPCs expressed canonical lymphatic markers (PDPN, LYVE1, VEGFR3 and F-actin) at 10 days of culture (fig. 6).

**Fig. 5.**
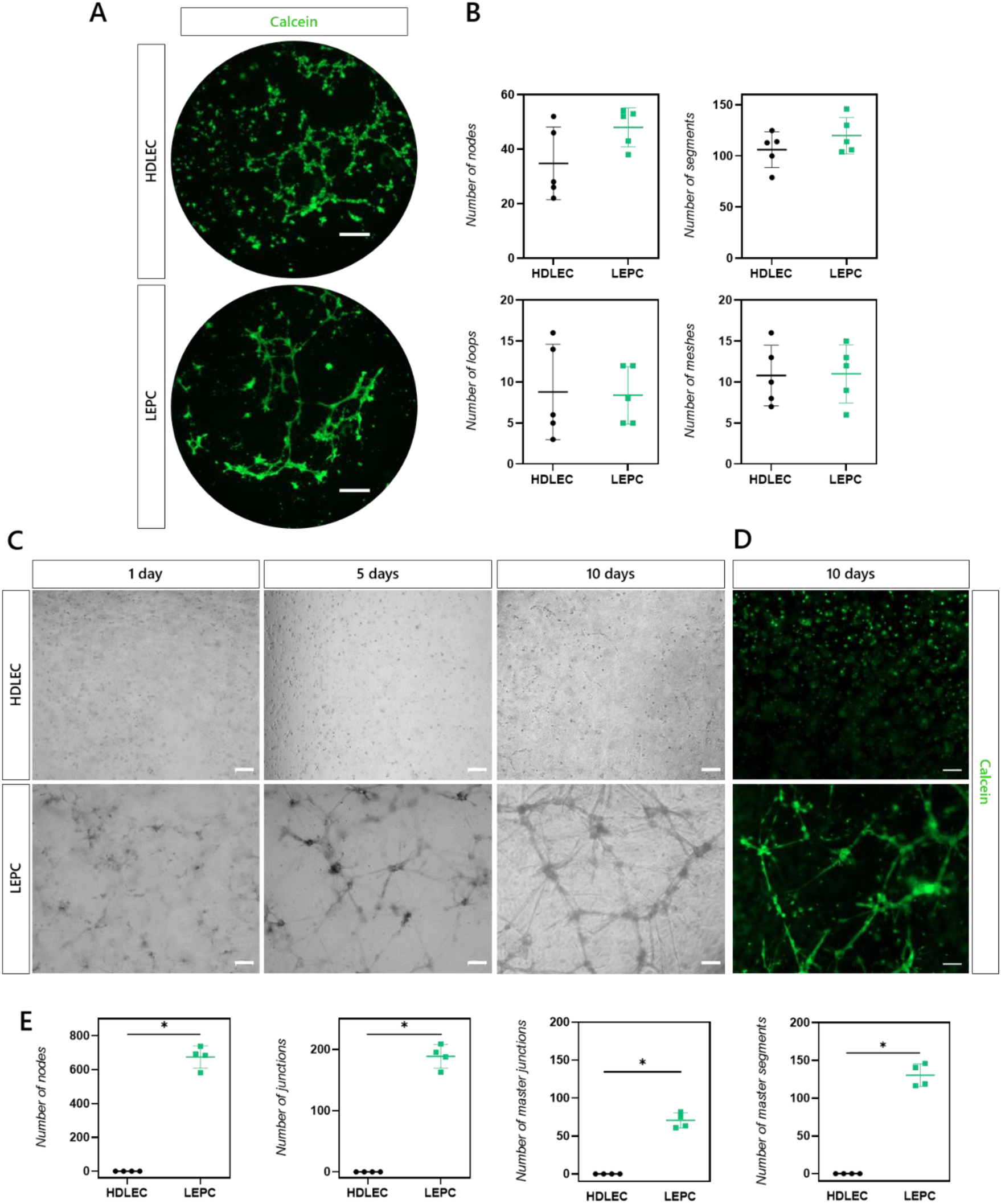
Stromal vascular fraction (SVF)-derived lymphatic endothelial progenitor cells (LEPCs) show better *in vitro* angiogenesis phenotype than human dermal lymphatic endothelial cells (HDLEC). (A) LIVE/DEAD staining images of SVF-derived LEPCs and HDLECs culture on a Matrigel (3 mg/mL) coating for 5 hours (2D angiogenesis assay). (B) Angiogenesis analysis of SVF-derived LEPCs compared to HDLECs in a 2D environment. N=5. (C) Bright field images of SVF-derived LEPCs and HDLECs embedded in a 3D Matrigel hydrogel (3 mg/mL) and cultured for 10 days. (D) LIVE/DEAD staining images of SVF-derived LEPCs and HDLECs embedded in a 3D Matrigel hydrogel (3 mg/mL) and cultured for 10 days. (E) Angiogenesis analysis of SVF-derived LEPCs compared to HDLECs in a 3D environment. n=4. Scale bars: (A) 500 µm; (C-D) 200 µm. Statistical significance: \**p*-value<0.05. Mann-Whitney test.

**Fig. 6.**
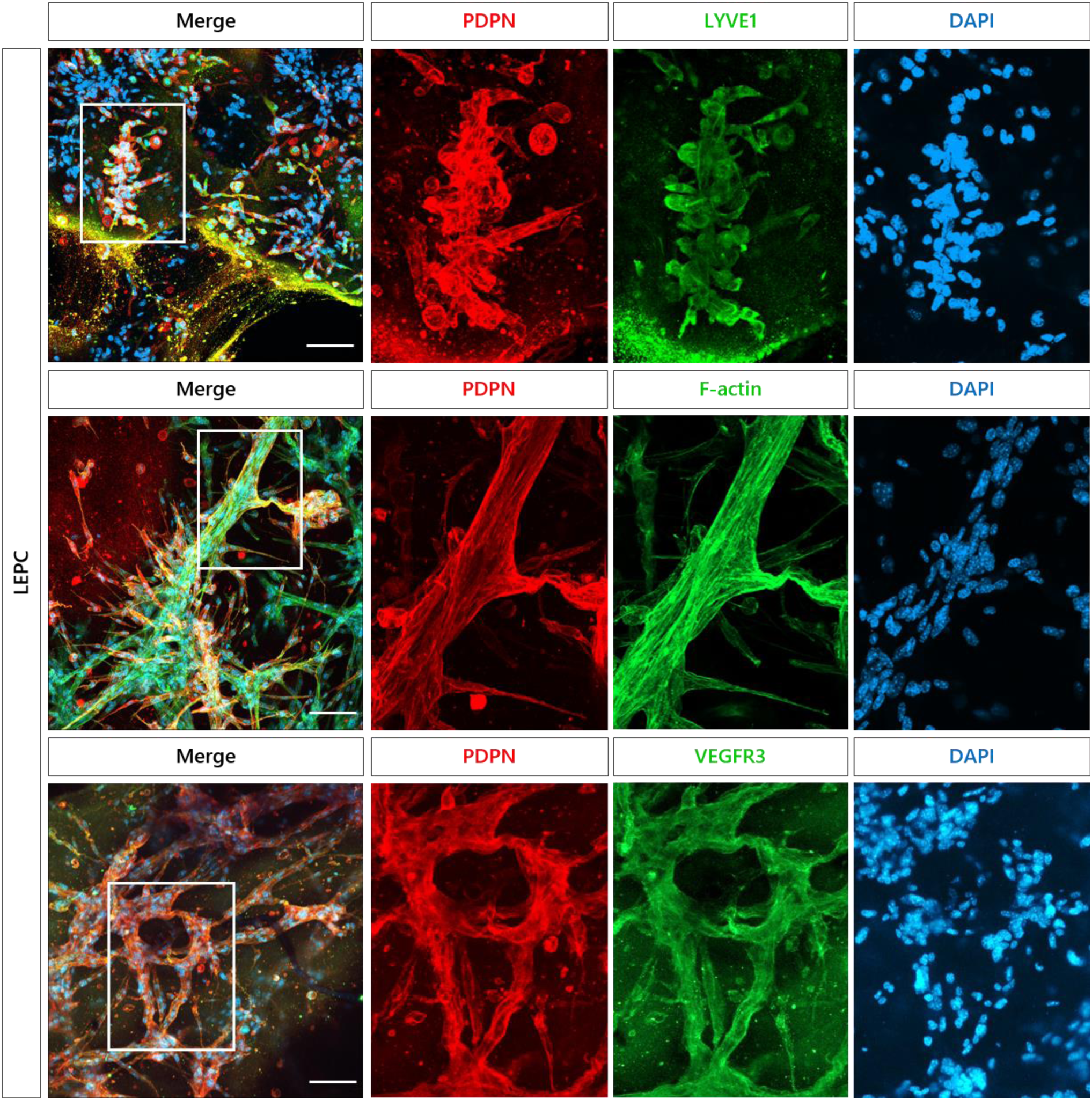
Stromal vascular fraction (SVF)-derived lymphatic endothelial progenitor cells (LEPCs) express canonical lymphatic markers in a 3D environment. Representative immunofluorescence staining images of LEPCs expressing lymphatic markers (PDPN, LYVE1, VEGFR3 and F-actin) embedded in Matrigel for 10 days. To the right, magnification of the white square in the left. Scale bar: 100 µm.

### RNA Sequencing confirms LEC lineage of SVF-derived LEPCs

For RNA sequencing, we utilized matched RNA samples isolated from SVF-derived LEPCs and their progenitor cells (passage 3). Three independent biological replicates were processed, with two technical replicates per sample. All cells underwent lymphatic differentiation prior to RNA extraction using TRI reagent solution, ensuring consistent sample origin across both genomic and expression analyses. Principal Component Analysis (PCA) of variance-stabilized transformed counts was performed to assess the overall variability and clustering of samples based on gene expression profiles. The first two principal components, PC1 and PC2, explained 69.2% and 9.3% of the total variance, respectively (fig. 7A). The PCA showed distinct clustering of the samples based on their experimental condition, also showing significant clustering in each experimental condition by their duplicates (fig. 7A). Consistent with the PCA, the sample clustering dendogram of the top 500 genes confirmed clear separation between biological groups, while duplicates exhibited high concordance (fig. 7B). Differential expression analysis showed a total of 2.037 upregulated and 1.936 downregulated genes for the differentiated cells in comparison to the MSCs (|log2 fold change| ≥1, adjusted *p*-value ≤0.05) (fig. 7C). Among the upregulated genes, several canonical markers of LECs were identified, including *Pdpn*, *Lyve1*, and *Vegfr3*, among others (fig. 7D). Focusing on the top 50 LEC specific genes, the differentiated cells showed a significant upregulation on 7 genes (*Pdpn, Vegfr3, Lyve1, Bmp2, Notch1, Cd34, Flt1*) as compared to their controls (fig. 7E). Additional details regarding the expression levels of these genes are shown in boxplots (comparing LEPCs and MSCs) in Sup. 2.

**Fig 7.**
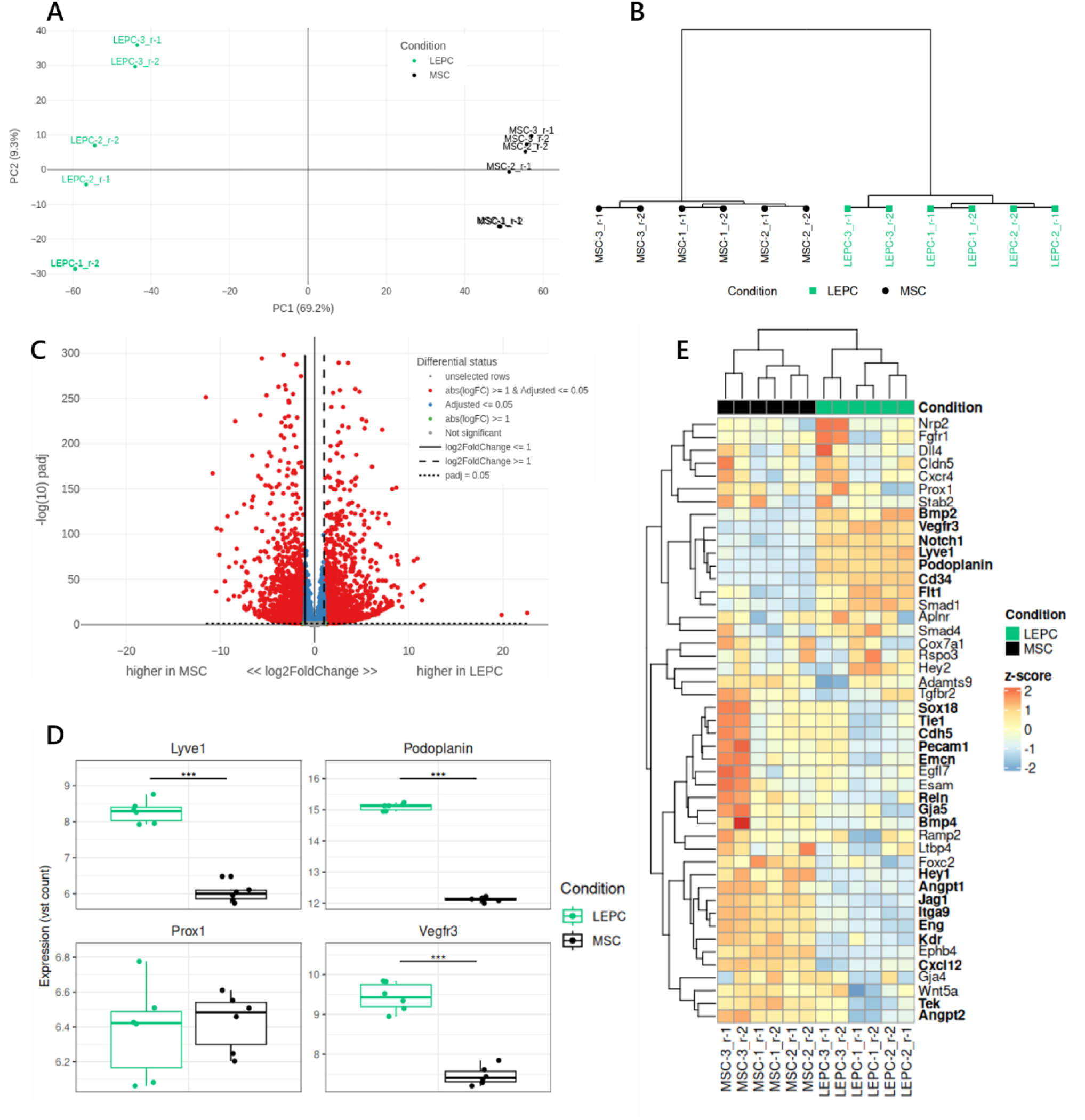
RNA sequencing analysis of Stromal vascular fraction (SVF)-derived lymphatic endothelial progenitor cells (LEPCs) confirm lymphatic genotype in comparison to mesenchymal progenitor cells. (A) Variance-stabilized transformed (VST) counts were used for Principal Component Analysis (PCA) plot. Axes show the top two principal components (PC1: 69.2%, PC2: 9.3%). Samples are colored by condition. (B) Sample clustering of the top 500 most variable genes. Distances between genes were estimated based on spearman correlation, which were then used to produce a clustering via the ward.D2 method. (C) Volcano plot of the differentially expressed genes (DEGs) between LEPCs and MSCs. Gray: non-significant; red: significant DEGs (|log2FC|≥1, adjusted *p*-value<0.05). (D) Boxplot of the differential expression of canonic lymphatic markers. Significance: ***adjusted *p*-value<0.001 (DESeq2 Wald test). (E) Heatmap of top 50 lymphatic endothelial cell (LEC) markers. Z-score-scaled VST counts for the 50 most significant LEC markers (rows) across samples (columns). Bold genes: |log2 fold change|≥1, adjusted *p*-value≤0.05. Clustering uses Euclidean distance and Ward.D2 linkage.

These findings demonstrate the potential to rapidly and precisely generate reliable lymphatic progenitors in vitro, which could outperform existing commercial cell lines for lymphatic-related applications.

## Discussion

Lymphatic tissue engineering has been significantly underexplored, resulting in a substantial knowledge gap that hampers the development of engineered constructs for purposes such as disease modelling, implants, or cell therapies. Tissue engineering typically relies on three essential components: cells, biomaterials, and biomolecules. However, in the context of lymphatic vessels/system, research has predominantly focused on biomolecules, as *in vivo* studies have proven valuable in understanding the aetiologies of various lymphatic system diseases.^2,49^ Among these, VEGF-C has received the most attention due to its critical role in lymphatic embryogenesis, postnatal stability, and lymphangiogenesis.^12,50,51^ Despite progress in understanding biochemical cues, much less attention has been given to the cellular aspects of lymphatic tissue engineering. Most studies involving cells have primarily examined their potential to regenerate dysfunctional or absent lymphatic vessels *in vivo.*^21^ These studies often use cells from a wide range of tissues, with a focus on MSCs or differentiated cells.^21^ For MSCs, the emphasis has typically been on their paracrine effects, which enhance lymphatic vessel formation, rather than on their direct differentiation and application.^25,52^

While *in vitro* studies have predominantly used commercial cell lines^53^, reliable protocols for differentiating MSC into LEPCs remain scarce^21^. Recognizing this significant gap, we have successfully established a novel *in vitro* differentiation protocol mimicking lymphatic embryogenesis, focusing on three clinically relevant tissue sources, obtainable from patients for autologous use. Among these, dermal tissue yielded the lowest cell count, whereas SVF and BM performed better. Even though, BM provides highest initial cell quantities, its poor attachment and proliferation in culture limits its utility. Conversely, SVF-derived MSCs, despite starting with fewer cells, proliferated quickly, achieving higher cell counts in half of the time, making them ideal for time-limited treatments^54^.

Both SVF- and BM-derived MSCs promote lymphangiogenesis through paracrine signalling^9,32,55^. However, adipose tissue is very highly related to the lymphatic pathologies, with a quite clear correlation of fat adipose tissue sedimentation and lymphatic malfunction^2^. Clinically, BM harvesting is highly invasive, posing greater risks to patients, while SVF extraction avoids this entirely^56–58^. In addition, SVF is easier to handle, requiring only standard culture conditions, unlike BM, which requires hypoxic conditions. Hence, SVF constitutes a superior option due its rapid expansion, minimal invasiveness and simplified culturing process.

Our novel differentiation protocol mimics the natural process of lymphatic embryogenesis and incorporates key molecular factors involved in adult lymphangiogenesis. VEGF-C is an early chemokine of the lymphatic embryogenesis (mice E9.5) that initiates the lymphatic commitment of venous endothelial cells, and driving the budding and sprouting of lymph sacs from the cardinal vein via VEGFR-3 signalling.^12,51,59^ CCBE1 acts synergistically to enhance VEGF-C activity through proteolytic maturation by ADAMTS3.^60^ Ang2 and 1 contribute to vascular remodelling by promoting vessel enlargement, branching and stabilizing the nascent vessels through TIE2 receptor.^50,61,62^ IGF2 plays a supplementary role during by activating VEGFR3-independet pathways, such as Akt and ERK, ensuring lymphangiogenesis continues even in the absence of VEGF-C.^63,64^ Finally, FGF2 promotes LEC tip cell formation, enabling vessel sprouting and elongation while stabilizing LECs via LYVE1 interaction.^65–67^ Together, these factors orchestrate the progression from initial LEC specification to the stabilization and maturation of a functional lymphatic network. Our results following this differentiation protocol align perfectly with the above-mentioned *in vivo* embryogenesis stages, as our denominated differentiation stages (induction, differentiation and maturation) accurately resemble embryogenesis *in vitro*. Our RT-qPCR results show that in comparison with the MSCs, the SVF-derived LEPCs show significant upregulation of the key lymphatic factors by statistically significant changes for *Pdpn*, *Lyve1*, *Vegfr3*, *Vegf-C* and *Igf2*. Based on the previous mentioned factors, the key LEC markers have always been PDPN, LYVE1 and VEGFR3. Overexpression of PDPN, LYVE1 and VEGFR3 is a hallmark of LEC identity and is pivotal for validating successful *in vitro* differentiation protocols.^68,69^ Furthermore, the overexpression of other markers, such as VEGF-C and IGF2, indicates that the cells not only express the key molecular biomarkers associated with LECs but also demonstrate the expression of functional markers. The overexpression of VEGF-C sustains the LEC phenotype by VEGFR3 signalling.^12,70,71^ Similarly, IGF2 can maintain lymphangiogenesis in a VEGF-C independent manner, suggesting that LEPCs have the capacity to self-sustain their lymphatic genotype even in the absence of VEGF-C.^64^

Following the successful differentiation of SVF-derived LEPCs, we performed extensive characterization to assess their behaviour and confirm their LEC phenotype *in vitro*. Our scratch-wound assay demonstrated a highly migratory phenotype comparable to HDLECs, as the SVF-derived LEPCs effectively closed the wound within 48 hours, displaying not only molecular marker expression but also robust functional capabilities. Additionally, maintaining the desired phenotype and gene expression over passages post-differentiation is crucial for reducing costs and accelerating practical applications.^72^ We observed that the expression of key markers, including *Pdpn*, *Lyve1*, *Vegfr-3*, and *Igf-2*, remained stable across the first two passages post-differentiation, indicating the robustness of our protocol. This stability could be attributed to the secretion of VEGF-C and IGF-2 by the cells, potentially forming an autocrine positive feedback loop.^12,64^ Furthermore, we assessed the ability of the cells to form angiogenic structures in both 2D and 3D environments. In 2D cultures, SVF-derived LEPCs readily formed web-like structures within 5 hours of Matrigel culture, mirroring angiogenesis assays and showing no significant difference compared to HDLECs. In 3D environments, our differentiated cells outperformed HDLECs, forming intricate tubular structures that simulated a lymphatic capillary network and expressing key lymphatic markers such as PDPN, LYVE1 and VEGFR3. These findings underscore the strength of our differentiation protocol, demonstrating the ability to derive functional LEPCs from SVF-derived MSCs *in vitro*, a breakthrough that holds promise for advancing disease modelling, tissue engineering, and cell therapy. For instance, disease modelling can drive therapeutic development for lymphedema and other traditional lymphatic diseases, as well as emerging conditions like obesity, where lymphatic dysfunction contributes to fat accumulation and inflammation,^73–75^ and Alzheimer’s disease, where the glymphatic system may play a pivotal role in addressing amyloid and tau pathology.^2^

Finally, the resulting transcriptomic analysis from the differentiation protocol provides very compelling evidence of successful differentiation, as demonstrated by the gene expression profiles, PCA clustering patterns, and differential expression analysis. Indeed, PCA results indicate that the first two principal components, PC1 (62.9%) and PC2 (9.3%), account for a large amount of the total variance. Such high levels of variance for PC1 suggests that the largest source of variability are the experimental conditions, i.e. the differentiation from MSCs into LEPCs. Furthermore, the significant clustering of the samples based on their experimental condition highlights the robustness and reproducibility of the differentiation protocol, which further supports the conclusion that the differentiation process induces significant changes in gene expression patterns that translate into phenotypical changes. The differential expression analysis revealed a very significant number of upregulated and downregulated genes in the differentiated cells compared to their mesenchymal progenitors, suggesting correct differentiation. When analysing the canonical genes for LECs, all genes confirmed our RT-qPCR results, indicating that there is a substantial upregulation of *Pdpn*, *Lyve1* and *Vegfr3*. The upregulation of these markers strongly suggests that the differentiated cells have acquired a LEPC phenotype. In addition, confirmation of the upregulation of most of the LEC specific markers, further confirms the correct differentiation of the mesenchymal stem cells into LEPCs. Collectively, our findings present a novel and previously unreported differentiation protocol for generating LEPCs in vitro, addressing a critical gap in lymphatic research. Additionally, we identify the optimal tissue source for LEPC isolation—a clinically accessible and highly relevant tissue with direct translational potential.

## Conclusion

This study explored the potential to differentiate MSCs derived from three distinct tissues into LEPCs for applications in lymphatic tissue engineering and cellular therapy. The findings demonstrated that SVF and BM serve as more reliable sources for isolating MSCs than the dermis. Furthermore, SVF-derived MSCs exhibited superior proliferation potential compared to BM-derived MSCs. A novel differentiation protocol was developed, mimicking the molecular signals of lymphatic embryogenesis, successfully guiding MSCs into LEPCs. Our results indicated that SVF-derived MSCs achieved more efficient differentiation into LEPCs than BM-derived MSCs, as evidenced by flow cytometry analysis of PDPN expression. Additional characterization of SVF-derived LEPCs confirmed their LEC phenotype, displaying high migratory capacity, a strong propensity for angiogenic structure formation, and consistent expression of key lymphatic markers in both 2D and 3D environments. Future studies will focus on evaluating these cells in *in vivo* models to assess their lymphangiogenic capabilities as a preliminary step toward their application in cell therapy. Overall, this research highlights a novel differentiation protocol that replicates the embryonic development of the lymphatic system, enabling the transformation of MSCs into LEPCs. These cells hold significant promise for tissue engineering, disease modelling, and therapeutic applications.

---

**Table S1.** Antibodies and other stainings used for immunofluorescence staining.

| Antigen | Type | Conjugated | Reference | Host | Reactivity | Fluorophore | Dilution | Brand |
| --- | --- | --- | --- | --- | --- | --- | --- | --- |
| PDPN | Primary | No | 127402 | Syrian Hamster | Mouse | No | 1/100 | BioLegend |
| LYVE1 | Primary | No | 103-PA50AG | Rabbit | Mouse | No | 1/100 | Reliatech |
| VEGFR3 | Primary | No | AF743 | Goat | Mouse | No | 1/100 | RD Systems |
| - | Secondary | Yes | A21451 | Goat | Hamster | Alexa Fluor 647 | 1/500 | Invitrogen |
| - | Secondary | Yes | A31572 | Donkey | Rabbit | Alexa Fluor 555 | 1/500 | Molecular Probes |
| - | Secondary | Yes | A11055 | Donkey | Goat | Alexa Fluor 488 | 1/500 | Molecular Probes |
| F-Actin | Phalloidin Staining | Yes | A12379 | - | - | Alexa Fluor 488 | 1/400 | Invitrogen |
| Nuclei | DAPI Staining | No | D3571 | - | - | DAPI | 1/1000 | Invitrogen |

**Table S2.** Antibodies used for flow cytometry analysis.

| Antigen | Type | Conjugate | Fluorophore | Reference | Host | Reactivity | Clone | Isotype | Dilution | Brand |
| --- | --- | --- | --- | --- | --- | --- | --- | --- | --- | --- |
| PDPN | Primary | Yes | APC | 127410 | Syrian Hamster | Mouse | 8.1.1 | IgG | 1/200 | BioLegend |
| SCA1 | Primary | Yes | PE-Cy7 | 558162 | Rat | Mouse | D7 | IgG2a<sub>k</sub> | 1/100 | BD |
| CD90.2 | Primary | No | - | 553009 | Rat | Mouse | 30-H12 | IgG2b | 1/200 | BD |
| CD105 | Primary | No | - | 14-1051-82 | Rat | Mouse | MJ7/18 | IgG2a<sub>k</sub> | 1/100 | Thermo Fisher |
| CD44 | Primary | Yes | PE | 553134 | Rat | Mouse | IM7 | IgG2b | 1/40 | BD |
| CD31 | Primary | Yes | PE | 553373 | Rat | Mouse | MEC13.3 | IgG2a<sub>k</sub> | 1/100 | BD |
| CD34 | Primary | Yes | PE | 551387 | Rat | Mouse | RAM34 | IgG2a<sub>k</sub> | 1/100 | BD |
| - | Secondary | Yes | Alexa Fluor 488 | A11006 | Goat | Rat | - | IgG | 1/500 | Molecular Probes |
| - | Isotype control | Yes | APC | 402012 | Syrian Hamster | Mouse | SHG-1 | IgG | 1/200 | BioLegend |
| - | Isotype control | Yes | PE | 553930 | Rat | Mouse | R35-95 | IgG2a<sub>k</sub> | 1/40 | BD |

**Table S3.**
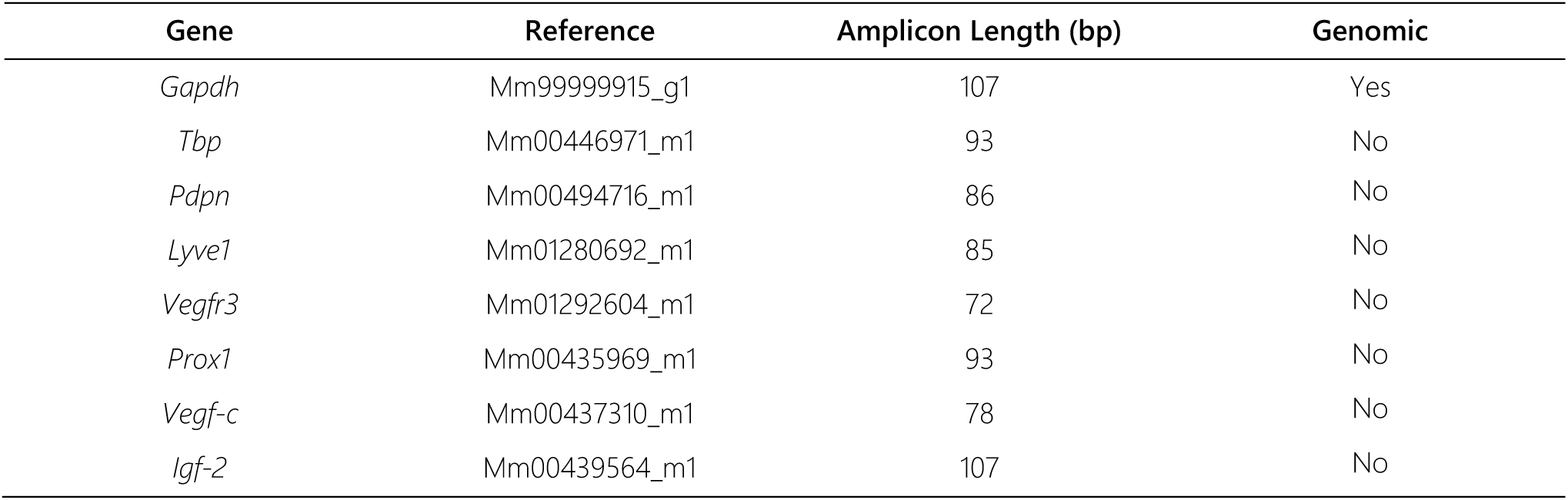
Taqman probes use for RT-qPCR analysis.

**Supplementary 1.**
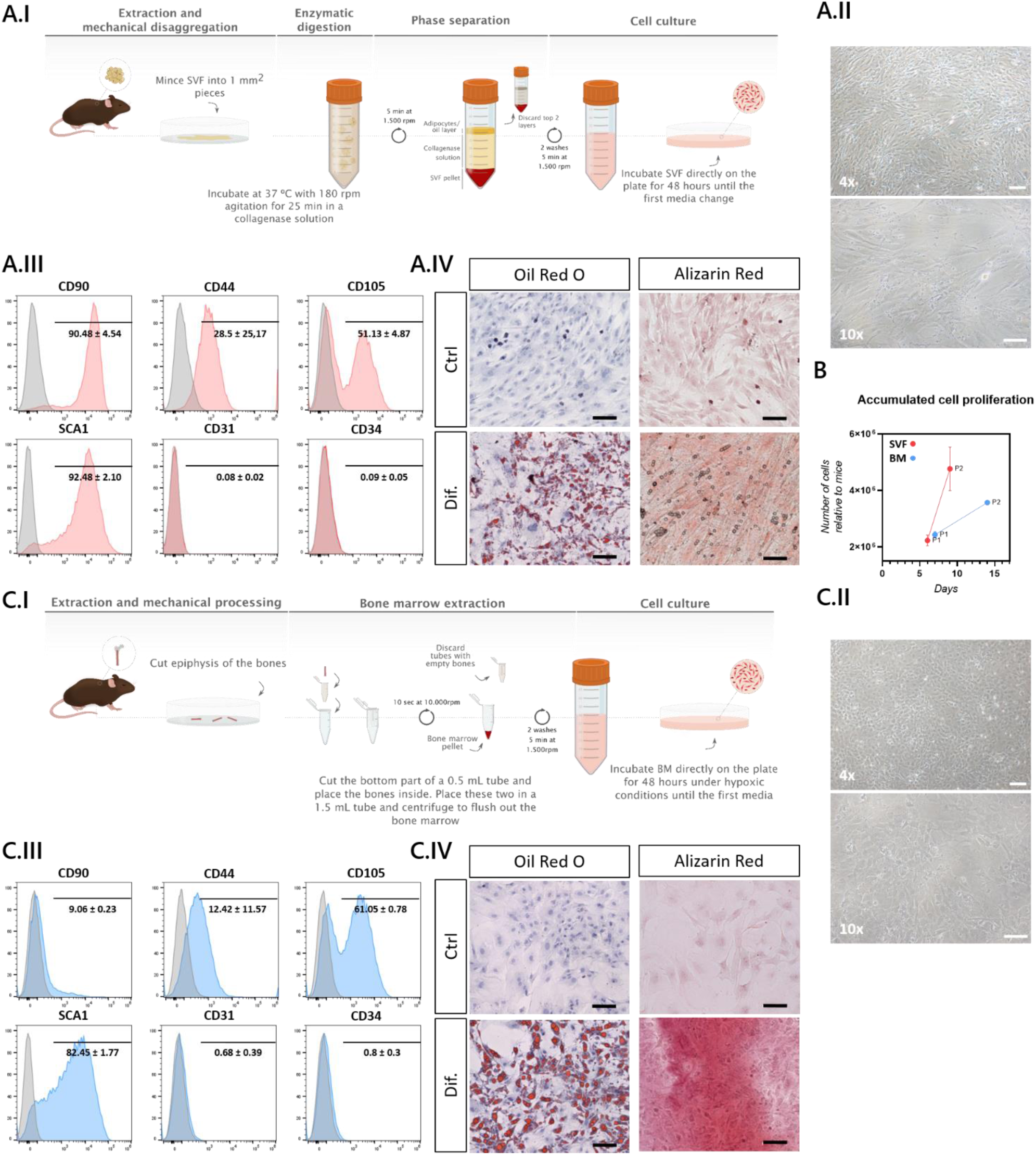
Mesenchymal stem cells (MSCs) isolation protocol and lineage confirmation. (A) Isolation and confirmation of stromal vascular fraction (SVF)-derived MSCs; (A.I) Illustration of the isolation protocol followed for obtaining SVF-derived MSCs; (A.II) Morphology of the SVF-derived MSCs in culture; (A.III) Expression of the canonical MSC markers by flow cytometry; (A.IV) Adipogenic differentiation of SVF-MSCs; (A.V) Osteogenic differentiation of SVF-MSCs. (B) Accumulated cell proliferation of SVF- and BM-derived MSCs over passages relative to mice. (C) Isolation and confirmation of BM-derived MSCs; (C.I) Illustration of the isolation protocol followed for obtaining BM-derived MSCs; (C.II) Morphology of the BM-derived MSCs in culture; (C.III) Expression of the canonical MSC markers by flow cytometry; (C.IV) Adipogenic differentiation of BM-MSCs; (C.V) Osteogenic differentiation of BM-MSCs.

**Supplementary 2.**
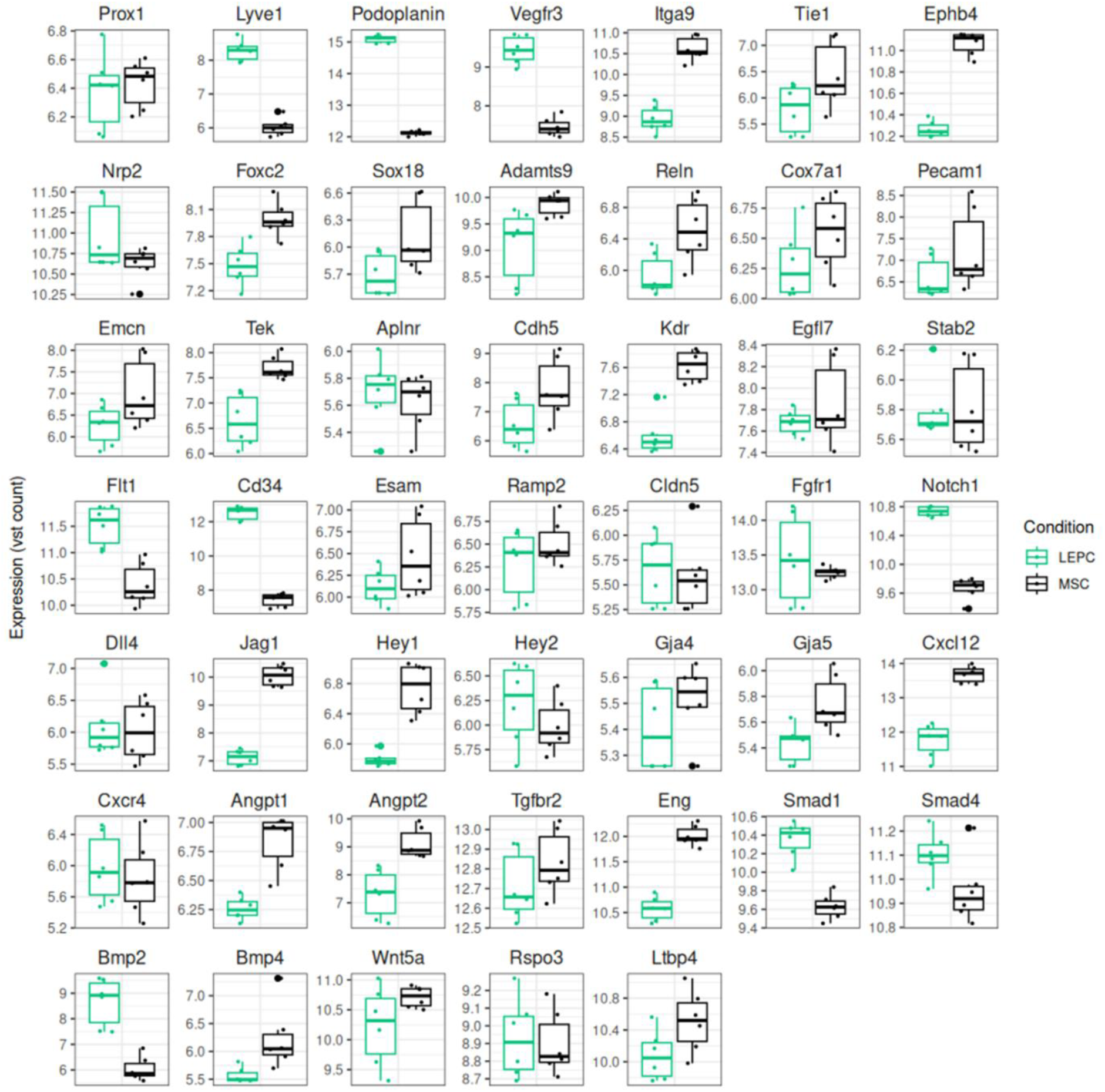
Boxplots of the top 50 lymphatic endothelial cell (LEC) markers expressed in lymphatic endothelial progenitor cells (LEPCs) in comparison to their mesenchymal progenitors.

